# Passive clearing and 3D lightsheet imaging of the intact and injured spinal cord in mice

**DOI:** 10.1101/2021.03.22.434816

**Authors:** Dylan A. McCreedy, Frank L. Jalufka, Madison E. Platt, Sun Won Min, Megan A. Kirchhoff, Ron Manlapaz, Eszter Mihaly, Jessica C. Butts, Nisha R. Iyer, Shelly E. Sakiyama-Elbert, Steven A. Crone, Todd C. McDevitt

## Abstract

The spinal cord contains a diverse array of sensory and motor circuits that are essential for normal function. Spinal cord injury (SCI) permanently disrupts neural circuits through initial mechanical damage, as well as a cascade of secondary injury events that further expand the spinal cord lesion, resulting in permanent paralysis. Tissue clearing and 3D imaging have recently emerged as promising techniques to improve our understanding of the complex neural circuitry of the spinal cord and the changes that result from damage due to SCI. However, the application of this technology for studying the intact and injured spinal cord remains limited. Here we optimized the passive CLARITY technique (PACT) to obtain gentle and efficient clearing of the murine spinal cord without the need for specialized equipment. We demonstrate that PACT clearing enables 3D imaging of multiple fluorescent labels in the spinal cord to assess molecularly defined neuronal populations, acute inflammation, long-term tissue damage, and cell transplantation. Collectively, these procedures provide a framework for expanding the utility of tissue clearing to enhance the study of spinal cord neural circuits, as well as cellular- and tissue-level changes that occur following SCI.

## 1 Introduction

The spinal cord mediates a complex repertoire of sensory and motor functions through a sophisticated network of neural circuits. Spinal cord injury (SCI) disrupts the transmission of neural signals and triggers widespread cellular and molecular events that mediate a secondary wave of tissue damage. While many different therapeutic strategies have been developed to limit secondary injury or promote repair of damage neural circuits (Griffin and Bradke 2020; Kwon, Okon, Hillyer, et al. 2011; Kwon, Okon, Plunet, et al. 2011; Tetzlaff et al. 2011), limited efficacy in the clinical setting has been achieved to date. Greater anatomical characterization of the intact and injured spinal cord may provide critical insight that can improve future studies to promote recovery after SCI. One of the major limitations to studying the spinal cord neural circuits, as well as the secondary pathological mechanisms that exacerbate tissue damage after SCI, has been the traditional use of labor-intensive and error-prone histological sectioning (Erturk et al. 2011; Steward, Zheng, and Tessier-Lavigne 2003).

Tissue clearing has emerged as a promising approach to render the opaque tissues of the central nervous system (CNS) optically transparent thereby enabling whole volumetric imaging of the entire tissue without the need for physical sectioning (Ueda et al. 2020; Richardson and Lichtman 2015). Many different tissue clearing protocols have been developed to date including hydrophobic clearing techniques that rely on organic solvents (e.g. 3DISCO), as well as hydrophilic and hydrogel-based techniques that utilize aqueous-based reagents (e.g. CUBIC and CLARITY, respectively) (Tainaka et al. 2018; Chung et al. 2013; Susaki and Ueda 2016; Erturk et al. 2012). Hydrophobic-based clearing techniques promote rapid and thorough clearing of CNS tissue, but can quench fluorescent protein signals and may impact endogenous epitopes needed for immunofluorescent (IF) labeling (Renier et al. 2014; Ueda et al. 2020). Hydrogel-based techniques can help preserve biomolecules, but often require additional forces such as electrophoresis to improve the speed and quality of tissue clearing (Chung et al. 2013; Tomer et al. 2014; Kim et al. 2015). Passive hydrogel-based protocols, such as the passive CLARITY technique (PACT), eliminate the need for specialized clearing equipment through reduced hydrogel crosslinking, which improves the rate of clearing and IF labeling while still maintaining sample integrity and transgenic fluorescence (Yang et al. 2014; Treweek et al. 2015).

Advances in tissue clearing techniques have enabled greater detailed analysis of intact neural circuits in the mouse brain, however, corresponding studies of the spinal cord have been more limited. Recent studies have demonstrated successful clearing of the murine spinal cord using hydrophobic clearing techniques, which has enabled the visualization of supraspinal, propriospinal and sensory circuits labeled with viral and non-viral tracers (Soderblom et al. 2015; Ni et al. 2014; Wang et al. 2018; Hilton et al. 2019; Pocratsky et al. 2017). Similar methods, albeit with the CLARITY technique, have been used to study reorganization of the cortico-reticulo-spinal pathway after SCI (Asboth et al. 2018). Thus, while promising, tissue clearing and 3D imaging techniques for the spinal cord remain under-utilized as a common form of analysis. The optimization and validation of a simple clearing method for the intact and injured spinal cord could enhance the use of tissue clearing and 3D imaging for studying spinal cord neural circuits and secondary injury after SCI.

In this study, we optimized the PACT tissue clearing procedure to obtain gentle and efficient clearing of the murine spinal cord for 3D imaging by light-sheet microscopy. We demonstrate successful IF labeling and preservation of transgenic fluorescent proteins in cleared spinal cords, which enabled the visualization and quantification of multiple neuronal subtypes. We also validated our clearing procedures in the injured spinal cord to assess acute inflammation and long-term white matter loss. Finally, we show IF labeling for transplanted human induced pluripotent stem cell (hiPSC)-derived interneurons that enabled the complete visualization of transplant cell survival and axon extension in the intact and injured spinal cord. Collectively, these results show the promising utility of PACT tissue clearing for studying the intact and injured spinal cord.

## 2 Materials and Equipment

### 2.1 Mouse Transcardial Perfusion and Dissection

1. Perfusion pump
2. 32% paraformaldehyde (Electron Microscopy Sciences, 15714-S)
3. 10X phosphate-buffered saline (Corning®, 20-031-CV)
4. 15 mL conicals (Falcon™, 352196)
5. 50 mL conicals (Falcon™, 352070)
6. Surgical tools — fine tweezers (FST, 91115-10), adson forceps (FST, 91106-12), Metzenbaum scissors (FST, 14016-14), micro-scissors (FST, 15025-10)

### 2.2 Hydrogel Polymerization

1. 15 mL conicals (Falcon™, 352196)
2. 40% Acrylamide (Calbiochem, 1150-OP) (3)
3. VA-044 (FujiFilm, 011-19365)
4. Parafilm (Parafilm™, PM996)
5. Nutating rocker — LabDoctor™ Mini Nutating Rocker (Ward’s Science™, H3D1020)
6. Incubator — Ward’s® Mini Digital Incubator (Ward’s Science™, 470230-608)
7. Vacuum desiccator (SP Scienceware™, F42025-0000)
8. Compressed nitrogen
9. Tube Revolver (Thermo Scientific™, 88881001)

### 2.3 Delipidation (Clearing)

1. Stirring Hotplate (Thermo Scientific™, SP88854100)
2. Boric Acid (Fisher Chemical™, A73-500)
3. Sodium Hydroxide (Millipore Sigma, S8045)
4. Sodium Dodecyl Sulfate (SDS) (Fisher BioReagents, BP82005)
5. Nutating rocker — LabDoctor™ Mini Nutating Rocker (Ward’s Science™, H3D1020)
6. Incubator — Ward’s® Mini Digital Incubator (Ward’s Science™, 470230-608)
7. Parafilm (Parafilm™, PM996)
8. Triton™ X-100 (Alfa Aesar, A16046-AE)
9. Tube Revolver (Thermo Scientific™, 88881001)
10. 15 mL conicals (Falcon™, 352196)

### 2.4 Immunohistochemistry (Optional)

1. Normal Donkey Serum (Millipore Sigma, D9663)
2. Sodium Azide (Millipore Sigma, S2002)
3. 0.22 μm Syringe Filter Unit (Millex-GP, SLGP033RS)
4. 10 mL syringe (Fisherbrand™, 14955458)
5. 1.5 mL microcentrifuge tubes (Denville, C2170)
6. 0.6 mL microcentrifuge tubes (Fisherbrand™, 05-408-120)
7. 15 mL conicals (Falcon™, 352196)
8. Primary antibodies — ChAT (1:200, Millipore Sigma, AB144P), NueN (1:200, Millipore Sigma, ABN78), Mouse S100A8 (1:500, R&D Systems™, AF3059), hTau (1:500, BioLegend®, MMS-520R)
9. Secondary antibodies (1:200) — Donkey anti-Goat IgG AlexaFluor® 488 (Invitrogen, A32814), Donkey anti-Mouse IgG AlexaFluor® 555 (Invitrogen, A32773), Donkey anti-Rabbit IgG AlexaFluor® 647 (Invitrogen, A32795)

### 2.5 Tissue Embedding and Refractive Index Equilibration

(10) Sodium phosphate, monobasic, monohydrate (Millipore Sigma, S9638)
(11) Sodium phosphate, dibasic, anhydrous (Millipore Sigma, 567550)
(12) Agarose, low-melting (Millipore Sigma, A9045)
(13) 0.5cc syringes (EXEL®, 26028)
(14) 15mm x 85 mm Culture Tubes (Pyrex, 99445-15)
(15) Nycodenz (Alere Technologies, 1002424)
(16) Heat block with 1.5 mL adaptor

## 3 Methods

### 3.1 Animals

Adult male and female C57Bl6, Chx10-Cre, Ai14 (B6.Cg-GT(ROSA)26Sor^tm14(CAG-tdTomato)Hze^/J), and NOD scid gamma (NSG; NOC.Cg-Prkdc^scid^Il2rg^tm1Wjl^/SzJ) mice were used for all studies. All studies were approved by the Institutional Animal Care and Use Committees at the University of California San Francisco, Texas A&M University, and the Cincinnati Children’s Hospital. All studies were in accordance with the United States Department of Agriculture guidelines.

### 3.2 Spinal cord injury

Adult (∼3-5 months) male and female mice were anesthetized with 2% isoflurane and a laminectomy was performed at the ninth thoracic vertebra (T9) to expose the spinal cord. A 70 kDyn spinal cord contusion was delivered using the Infinite Horizons impactor. The musculature was sutured closed and the skin closed with wound clips. Manual bladder expression was performed twice daily and subcutaneous injections of saline and antibiotics were given daily.

### 3.3 V2a interneuron transplantation

Human induced pluripotent stem cell (hiPSC)-derived V2a interneurons were generated as previously described (Butts et al. 2019). For transplantation into the intact spinal cord, adult female NSG mice were anesthetized with 2% isoflurane and a laminectomy was performed at T9. The spinal column was secured using the IH spinal clamps and approximately 500,000 hiPSC-derived V2a interneurons were transplanted over four injections sites into the ventral grey matter at T9 using a glass micropipette attached to a Nanoject III (Drummond). In another cohort of NSG mice, the T9 laminectomy site was re-exposed at 14 days post-SCI and approximately 500,000 hiPSC-derived V2a interneurons were transplanted in the ventral grey matter bilaterally at the rostral and caudal aspects of T9. Following transplantation, the musculature and skin were closed and postoperative care was performed as described above for spinal cord injury.

### 3.4 PACT tissue clearing and immunofluorescent labeling

#### Step 1: Mouse Transcardial Perfusion-Fixation

1. Prepare 1X PBS and 4% paraformaldehyde in 1X PBS (pH 7.4) solutions and place on ice. **Caution:** Paraformaldehyde is a carcinogenic agent. Ensure to take the proper handling precautions; perform all procedures in a fume hood. Paraformaldehyde waste should be discarded according to institutional protocols.
2. Perfuse mouse with 25 mL of ice-cold 1X phosphate buffered saline (PBS) followed by 25 mLs of ice-cold 4% paraformaldehyde in PBS.
3. Dissect out the spinal column using Metzenbaum scissors and adson forceps. Transfer the spinal column to 10 mL of ice-cold 4% paraformaldehyde in 1X PBS in a 15 mL conical and post-fix overnight (O/N) at 4°C.
4. The following day, transfer the spinal column to 40 mL of ice-cold 1X PBS in a 50 mL conical and store O/N at 4°C. **Checkpoint:** Possible stopping point – tissue may be stored in 1X PBS with 0.01% (w/v) sodium azide at 4°C for several weeks.

#### Step 2: Spinal Cord Dissection and Meninges Removal

1. Dissect the spinal cord from the spinal column.
  1.1 Starting at the cervical end of the cord, locate the interior of the spinal column and carefully make lateral incisions using micro-scissors on both sides of the ventral aspect of the first vertebrate. **Note:** Micro-scissors should be oriented flat along the ventral side of the vertebrate to prevent damage to the spinal cord.
  1.2 Using fine tweezers, peel off the cut portion of the vertebrae to expose the spinal cord. Repeat for each vertebrate down the entire length of the cord.
2. Remove spinal cord from spinal column.
  2.1 Starting at the lumbar end of the cord, sever the spinal nerves that anchor the cord in the column. Carefully peel away the spinal cord from the rest of the column.
  2.2 Remove hair, bone fragments, and debris adhered to the spinal cord.
3. Remove the meninges.
  3.1 Transfer the spinal cord to petri dish. Starting at either end of the spinal cord, carefully peel off the meninges using fine forceps while holding onto the spinal cord with adson forceps.
  **Note:** Occasionally wet the spinal cord with 1X PBS as needed to prevent tissue from drying out. Removal of meninges is a critical step towards faster and uniform clearing. Take special care not to tug on the meninges as such force will result in severing the tissue.
4. Transfer the dissected spinal cord to 14 mL of ice-cold 1x PBS in a 15 mL conical. **Checkpoint** : Possible stopping point – tissue may be stored in 1x PBS with 0.01% (w/v) sodium azide at 4°C for several weeks.

#### Step 3: Hydrogel Polymerization

1. Prepare 12.5 mL/sample of A4P0 (4% acrylamide + 0% paraformaldehyde) monomer solution.
  1.1 For 40 mL of solution, combine 4 mL of 10x PBS and 4 mL of 40% acrylamide in 32 mL of ice-cold ddH2O.
  1.2 Chill on ice for 30 minutes.
  1.3 Dissolve 100 mg of VA-044 in to the chilled 4% acrylamide in 1X PBS solution.
  **Caution**: Acrylamide is a carcinogenic agent. Always ensure proper handling precautions, work with acrylamide in a fume hood. Acrylamide waste should be discarded according to institutional protocols.
2. Transfer the spinal cord to 12.5 mL of ice-cold A4P0 monomer solution in a 15 mL conical.
3. Seal with parafilm around the cap of conical to prevent leakage. Secure conical horizontally to a tray that can contain any potential leaks.
4. Incubate O/N at 4°C with gentle rocking on a nutating rocker. **Note:** Place the conical flat on the nutating rocker such that the spinal cord tissue partially inverts during rocking. The movement of the spinal cord tissue during inversion will facilitate penetration of the A4P0 hydrogel.
5. Transfer the conical containing the spinal cord to a fume hood, remove parafilm, and uncap the conical. Place the uncapped conical upright in a tube rack within a vacuum desiccator and purge the spinal cord of residual oxygen for 30 min. Occasionally tap the desiccator to dislodge bubbles.
6. Flood the dessicator with nitrogen for 1-2 minutes and then cap the conical inside the desiccator. Seal the conical with parafilm.
7. Incubate the conical with the spinal cord tissue upright in pre-heated 37°C incubator for 3 hours.
8. Remove the sample from the polymerized hydrogel solution. **Note:** Solution should be viscous, but will still flow when the conical is inverted.
9. Wash the sample in 14 mL of 1x PBS in a new 15 mL conical on a tube revolver for 5 minutes at room temperature (RT).

#### Step 4: Delipidation (Clearing)

1. Prepare 1 M boric acid buffer (pH=8.5)
  1.1 Dissolve 61.83 g of boric acid and 12 g of sodium hydroxide pellets (NaOH) in 900 mL of ddH_2_O. Stir until fully dissolved and clear. Add ddH_2_O up to final volume of 1L
2. Make 10% sodium dodecyl sulfate (SDS) in water (w/v).
  2.1 Dissolve 100 g of SDS micropellets (Fisher BioReagents, BP82005) in 800 mL of ddH_2_O. Stir constantly over heat up to 60°C. When solution is clear, turn off heat and cool to RT. Add ddH_2_O up to final volume of 1L.
  **Caution**: Sodium dodecyl sulfate is a corrosive irritant. Always ensure the proper handling precautions; make solution and perform all solution changes in a fume hood. SDS waste should be discarded according to institutional protocols.
3. Prepare 12.5 mL/sample of 8% SDS in 0.2 M boric acid buffer (SDS clearing solution) by combining 10 mL of 10% SDS with 2.5 mL of 1 M boric acid buffer. **Note:** Do not store the SDS clearing solution for more than one day since SDS may precipitate in boric acid buffer at RT.
4. Transfer tissue from PBS to 12.5 mL of SDS clearing solution in a 15 mL conical. Seal lid with parafilm and incubate at 37°C for two days on a nutating rocker. **Note:** Place the conical flat on the nutating rocker such that the sample partially inverts and moves toward each end of the conical during rocking. The movement of the spinal cord tissue will facilitate removal of lipids.
5. Replace SDS clearing solution every other day until the spinal cord is optically transparent. Full-length mouse spinal cord tissue should clear in 5-7 days. **Note:** Avoid leaving spinal cord tissues in SDS clearing solution after becoming optically transparent, as this can lead to sample degradation.
6. Prepare 0.2M boric boric acid buffer with 0.1% Triton™ X-100 (BBT)
  6.1 Combine 800 mL of ddH_2_0, 200 mL of 1 M boric acid buffer, and 1 mL of Triton™ X-100. Stir at RT until all components are fully dissolved.
7. Once the spinal cord has finished clearing, transfer to 14 mL of BBT in a 15 mL conical and rotate at RT for 30 min on tube revolver to wash out the residual SDS.
8. After 30 min, replace the BBT solution and continue rotating.
9. Repeat Step 5 four times then leave the sample in BBT solution O/N rotating at RT. **Note:** Rapid washing of the SDS solution is critical to preventing precipitation of SDS inside the sample, which reduces optical clarity and efficiency of IF labeling. The sample may become slightly opaque after washing, but should remain transparent. **Note:** Samples with endogenous or transgenic fluorescent signals should be protected from light during delipidation and washing. **Checkpoint:** Possible stopping point – tissue may be stored in BBT or 0.2 M boric acid buffer with 0.01% sodium azide at 4°C.

#### Step 5: Immunofluorescent (IF) labeling

1. Prepare staining buffer: BBT + 2% normal donkey serum (NDS) + 0.01% sodium azide.
2. Prepare primary antibodies in staining buffer (∼1 mL/sample).
3. Transfer the spinal cord tissue into the staining buffer with primary antibodies in a 1.5 mL MCT. Protect from light and rotate for two days at RT on tube revolver. **Note:** For small spinal cord tissues, transfer sample to ∼500μL of staining buffer with primary antibodies in a 0.6 mL MCT. **Note:** Spinal cord tissues should invert during rotation on the tube revolver, which will improve the penetration of antibodies. Two days is usually sufficient for antibodies to diffuse to the center of the spinal cord samples. Steps 1-3 can be repeated if necessary.
4. Transfer spinal cord tissue to 14 mL of BBT solution with 0.01% (w/v) sodium azide in a 15 mL conical and wash tissue with 5 buffer changes within 3-5 hours on a tube revolver at RT. Leave the sample in the last wash O/N on a tube revolver at RT. **Note:** From this point on, it will be crucial to add sodium azide to a final concentration of at least 0.01% to every solution to prevent infection/contamination of the sample.
5. Prepare secondary antibodies in staining buffer (∼1.2 mL/sample).
6. Filter staining buffer with secondary antibodies using syringe filter.
7. Transfer spinal cord tissue to 1 mL of filtered antibody solution in a 1.5 mL MCT. Protect from light and rotate for two days at RT on tube revolver. **Note:** For smaller tissues, prepare ∼700 μL of staining buffer with secondary antibodies. Transfer smaller tissue to ∼500 μL of filtered antibody solution in a 0.6 mL MCT.
8. Repeat step 4. **Checkpoint:** Possible stopping point – spinal cord tissue may be stored in BBT with 0.01% sodium azide at 4°C protected from light.

#### Step 6: Tissue Embedding and Refractive Index Equilibration

1. Prepare 0.1 M phosphate buffer (PB), pH 7.4.
  1.1. Dissolving 3.1 g of sodium phosphate (NaH_2_PO_4_, monobasic, monohydrate) and 10.9 g of sodium phosphate (Na_2_HPO_4_, dibasic, anhydrous) in 900 mL of ddH_2_O. Add ddH_2_O up to final volume of 1L.
2. Prepare 0.02 M phosphate buffer (pH 7.4) by combining 200 mL of 0.1 M PB and 800 mL of ddH_2_O.
3. Prepare 1.5% agarose solution by dissolving 300 mg of low-melt agarose in 20 mL of 0.02 M PB.
  3.1. Microwave solution in 10 – 15 second intervals until agarose is fully dissolved.
  3.2. Allow solution to cool to approximately 50°C and aliquot 1 mL into 1.5 mL MCTs and place on a heat block set at 42°C.
  **Note:** Extra agarose aliquots may be stored at 4°C, re-melted on a heat block at 95°C, and cooled to 42°C.
4. Cut off the needle tip of a 0.5cc insulin syringe such that the full internal diameter of the syringe is exposed. Pull up approximately 250-300 uL of agarose solution into the syringe.
5. Wash the sample by briefly placing in the agarose solution, then moving to a new MCT with agarose solution. Draw up the sample + agarose solution into the second half of the syringe. The syringe should be nearly full of agarose solution with the sample located near the tip of the syringe. **Note:** Ensure that there are no air bubbles around the tissue or near the plunger that may hinder imaging or cause the sample to slip out of the syringe, respectively.
6. Allow agarose solution to solidify protected from light for a minimum of one hour at RT.
7. Prepare enough refractive index matching solution (RIMS) to equilibrate samples in 5 mL each and to fill the sample chamber for lightsheet imaging.
  7.1 For ∼ 40 mL of RIMS, dissolve 30 g of Nycodenz in 22.5 mL of 0.02 M PB in a 50 mL conical. Place on a tube revolver for several hours until fully dissolved.
  7.2 Add sodium azide to a final concentration of 0.01%.
  7.3 Adjust the refractive index of the RIMS with Nycodenz (for low RIs) or 0.02 PB (for high RIs) until the RI is between 1.455-1.458. Store at 4°C.
8. Place the syringe in 5 mL of RIMS in a 15mm x 85mm culture tube.
9. Center the syringe in the opening of the culture tube and use parafilm to secure in place such that the tip of the syringe is just below the surface of RIMS.
10. Eject the embedded sample into the RIMS so that it is surrounded by the RIMS with the remaining agarose still inside of the syringe.
11. Equilibrate sample for 1-2 days in the dark at RT. Sample is ready to image once the sample and agarose are optically transparent in the RIMS. **Note:** Mounted tissue may be washed with 0.2 M boric acid buffer for 1 hour, gently removed from the agarose, and re-stained/re-mounted or stored in 0.2 M boric acid buffer with 0.01% sodium azide at 4°C. **Checkpoint:** Possible stopping point – mounted tissues may be stored in RIMS with 0.01% sodium azide at 4°C.

### 3.5 DiD labeling

Spinal cord samples were incubated in 10mM SDS in PBS (SWITCH-off) overnight at 37°C with gentle rocking before incubating with DiD (5ug/mL) in SWITCH-off buffer overnight at 37°C with gentle rocking. Spinal cord samples were then washing in PBS with 0.01% triton (SWITCH-on) overnight at 37°C with gentle rocking.

### 3.6 Lightsheet imaging

Prior to imaging, spinal cords were mounted in 1-2% agarose inside of 0.5mL syringes. After solidification of the agarose, spinal cords were suspended in refractive index matching solution (RIMS; 30g of Nycodenz in 0.02M phosphate buffer with 0.01% sodium azide; RI = 1.458) for two days. After equilibration in RIMS, spinal cord samples were imaged using a Zeiss Z1 lightsheet microscope equipped with a 5x objective. Image Z-stacks were captured for both left and right sided lightsheet illumination using Zen software. Dual side fusion of left and ride sided images was performed in Zen software.

### 3.7 Image processing and analysis

Maximum intensity projections (MIPs) of image Z-stacks were generated using Zen software. Composite MIP images were stitched from individual MIPs in Photoshop (Adobe). Image Z-stacks were visualized in Imaris 3D software (Bitplane). 3D videos were generated using the animation tool in Imaris. Soma and neutrophil quantification were performed using the spot assay in Imaris.

## 4 Results

### 4.1 Rapid and efficient clearing of spinal cord tissue through optimization of PACT

Previous protocols for PACT describe a broad range of conditions to accommodate the clearing of many different tissues (Treweek et al. 2015; Yang et al. 2014). In this study, our goal was to optimize the conditions of the PACT procedure to enable reproducible clearing and IF labeling in the mouse spinal cord (Figure 1A). With the optimized PACT procedures (see Methods for details), whole mouse spinal cords were efficiently cleared in approximately six days (Figure 1B). Cleared spinal cords became fully transparent after two days of incubation in refractive index matching solution (RIMS). Since no specialized equipment is required, multiple samples can be cleared in parallel to achieve high throughput processing. Passive clearing using hydrophilic and hydrogel-based techniques often require multiple weeks to months to achieve complete transparency (Tomer et al. 2014; Zhang et al. 2014; Woo et al. 2016), thus the optimized PACT procedure provides a simple and rapid method to passively clear the mouse spinal cord.

**Figure 1.**
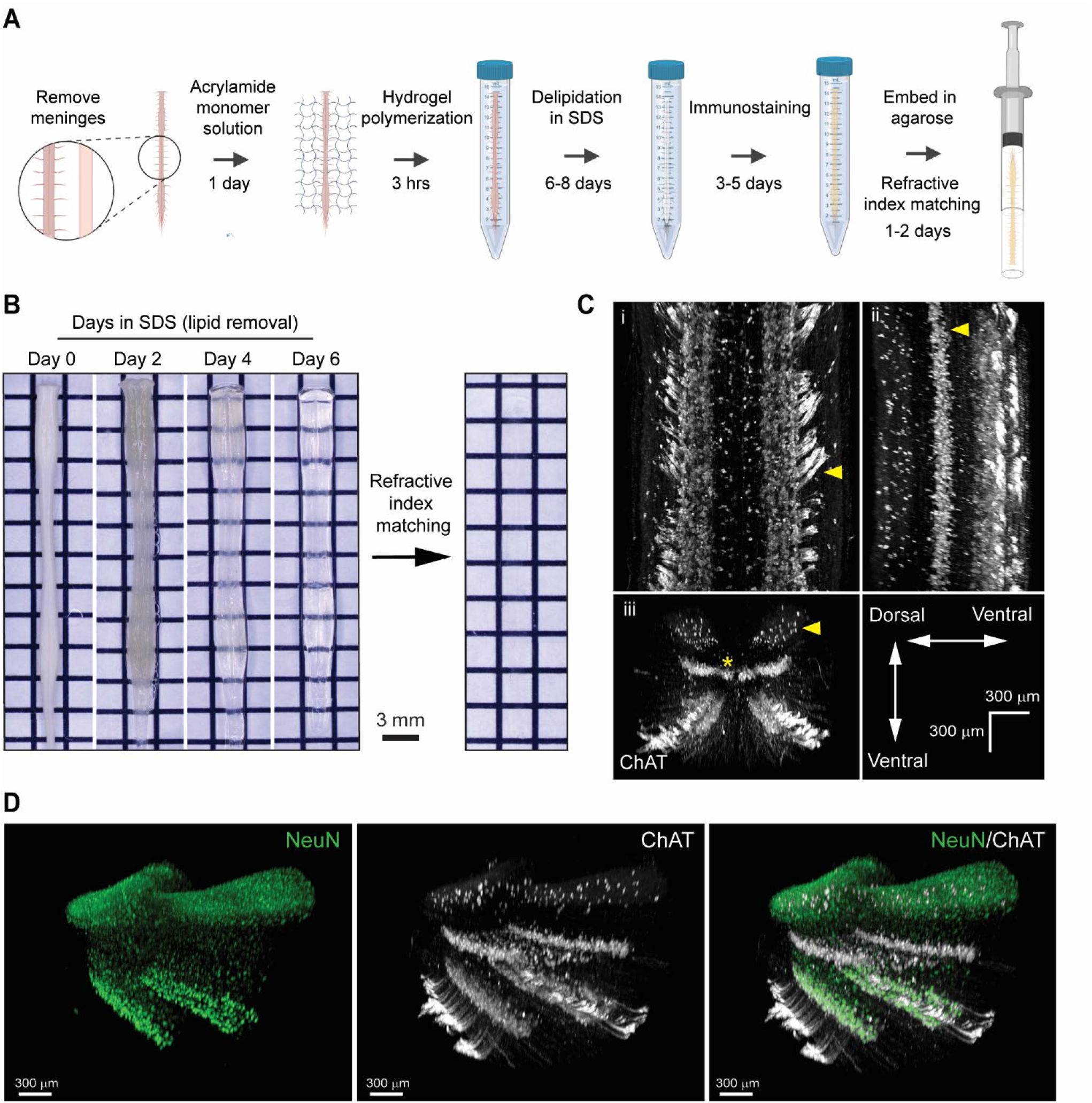
Optimization of PACT procedures enables efficient clearing and IF labeling of the murine spinal cord. A) Schematic of the optimized PACT clearing and IF labeling procedures. Created with BioRender. B) Comparison of spinal cord tissue clearing with the original (i) and optimized (ii) PACT procedures. C) IF labeling of ChAT with the original PACT protocol. D) 3D projections of ChAT IF labeling with the optimized PACT protocol in the thoracic mouse spinal cord from dorsal (i), sagittal (ii), and coronal (iii) views. Arrows indicate motor axons, cholinergic neurons in the intermediate zone, and dorsal cholinergic interneurons, respectively. Asterisk indicates putative V0c interneurons in the medial region of the intermediate zone. D) 3D projection of NeuN (green) and ChAT (white) IF labeling in the cleared thoracic mouse spinal cord. Abbreviations: choline acetyl transferase (ChAT), immunofluorescent (IF).

To determine the efficacy of IF labeling after PACT clearing, we incubated the cleared spinal cords with antibodies against ChAT and NeuN, followed by AlexaFluor-conjugated secondary antibodies, and imaged the samples using lightsheet microscopy. In the 3D images, robust labeling of several cholinergic (ChAT^+^) cell populations was observed (Figure 1C, Movie 1). Motor neurons were readily apparent in the ventral grey matter and groups of motor axons were seen extending away from the ventral horn towards the ventral roots (Figure 1Ci, arrow). Additional cholinergic populations were observed in the intermediate zone (Figure 1Cii, arrow), which may include a medial V0c interneuron population (Figure 1Ciii, asterisk) as well as a lateral population of motor neurons in the intermedial lateral nuclei. We also found a sparse cholinergic dorsal interneuron population (Figure 1Ciii, arrow), which has been previously described (Pawlowski et al. 2013). Specific IF labeling of NeuN was also observed, which enabled the assessment of NeuN and ChAT co-localization to confirm the grey matter location and neuronal identity of ChAT^+^ cells (Figure 1D). These data demonstrate bright IF labeling for easy detection of neuronal subclasses using cell-type specific markers in cleared spinal cord tissue.

### 4.2 PACT clearing preserves transgenic fluorescence for 3D imaging of spinal cord neural circuits

To determine the compatibility of the optimized PACT protocol with transgenic fluorescent protein signals in the murine spinal cord, we bred Chx10-Cre mice with the Ai14 reporter mice to generate offspring (Chx10::tdTomato mice) with tdTomato expression in V2a interneurons. Using lightsheet microscopy, we imaged the majority of the length of the cleared spinal cord (C1-L1) by stitching together multiple fields of view. Transgenic tdTomato fluorescent signal was well-preserved in V2a interneuron soma and axons in the cleared spinal cords of Chx10::tdTomato mice (Figure 2a, Movie 2). We observed tdTomato^+^ V2A interneuron soma distributed throughout the grey matter at all levels of the spinal cord, as well as extensive tdTomato^+^ axon tracts in the ventral spinal cord that spanned the length of the imaged region.

**Figure 2.**
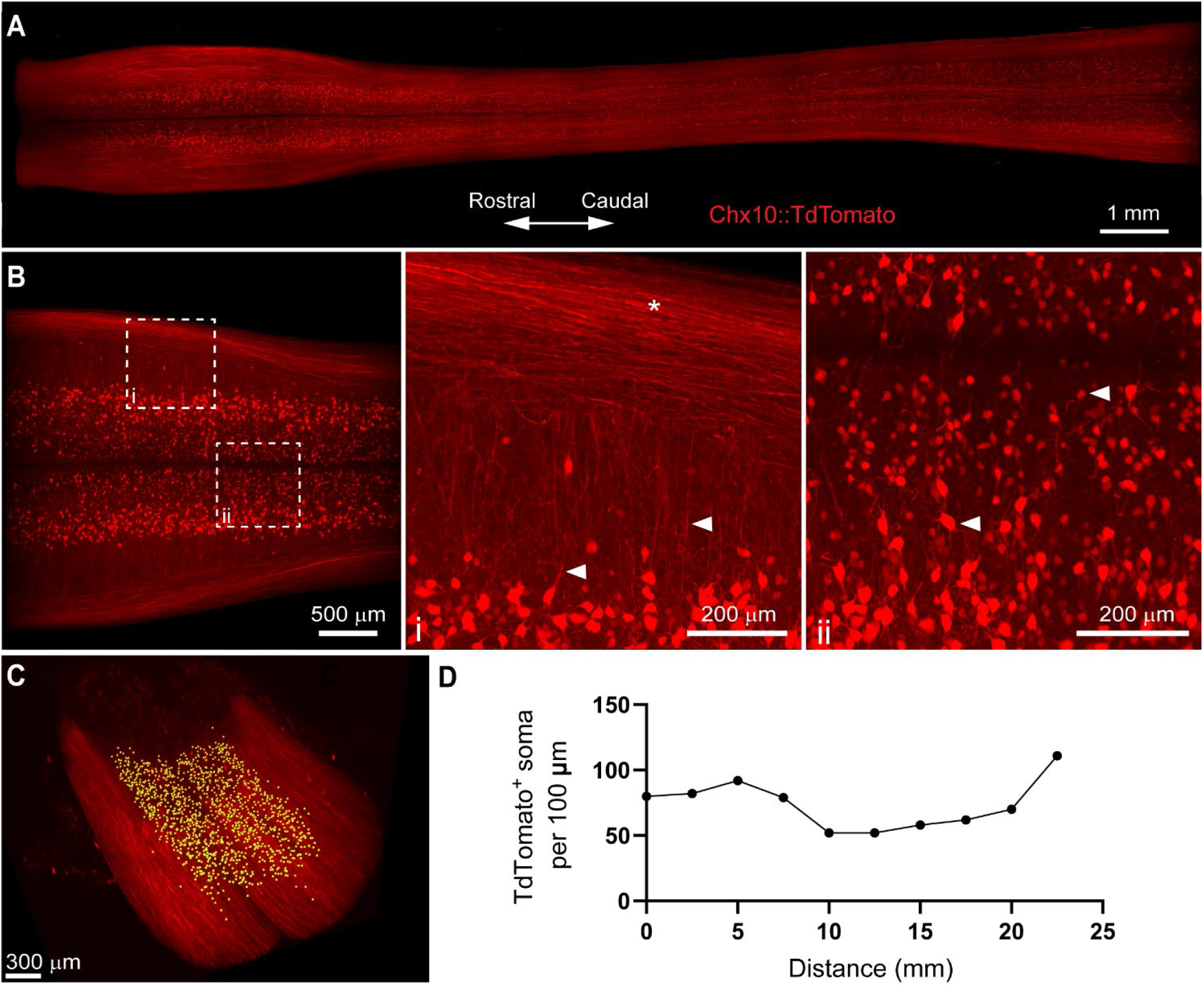
PACT clearing preserves endogenous tdTomato fluorescence in spinal cord interneurons for 3D imaging of neural circuits. A) Maximum intensity projection of tdTomato in V2a interneurons in the cleared spinal cord from Chx10::tdTomato mice. B) Maximum intensity projection of the intermediate zone from a single field of view of the cleared spinal cord showing V2a interneuron soma. Axons from V2a interneurons project laterally in the grey matter before extending along white matter tracts (i; arrows and asterisk, respectively). Large and small soma can be easily identified (ii, arrows) in the grey matter. C) V2a interneuron soma counting in Imaris (yellow dots). D) Average number of V2a interneurons per 100 um assess at every 2.5 mm along the length of the spinal cord.

To digitally isolate V2a interneurons for further analysis, we acquired maximum intensity projections of the z-stack image planes spanning the intermediate zone of the spinal cord including most of laminae VII (Figure 2B). We observed tdTomato^+^ axons from V2a interneurons extending laterally through the grey matter before joining tdTomato^+^ axon tracts in the spinal cord white matter (Figure 2Bi; arrows and asterisk, respectively). In addition, heterogeneity in tdTomato^+^ V2a interneuron soma size and location along the medialateral axis of the spinal cord was readily observed (Figure 2Bii, arrows). We quantified tdTomato^+^ soma numbers using Imaris software to assess V2a interneuron distribution along the length of the spinal cord (Figure 2C). V2a interneuron soma number appeared to vary along the length of the spinal cord, with putative increases in the cervical and lumbar regions (Figure 2D). This proof-of-concept experiment demonstrates how PACT tissue clearing and 3D imaging can be applied to transgenic tdTomato fluorescent protein expression to investigate molecularly-defined subclass of neurons in spinal cord neural circuits.

### 4.3 PACT clearing enables 3D imaging and quantification of acute inflammation after SCI

Inflammation after SCI has been predominantly characterized using tissue sections or by flow cytometry (Beck et al. 2010; Stirling and Yong 2008). While important, these techniques fail to fully capture spatial information that may provide insight into the dynamics of various immune cell populations within the spinal cord parenchyma. To determine if tissue clearing and lightsheet imaging could provide unique assessment of the distribution of immune cells after spinal cord injury, we performed a contusive SCI and isolated spinal cords the following day. Despite the noticeable hemorrhaging, PACT tissue clearing successfully rendered the acutely injured spinal cord optically transparent (Figure 3A).

**Figure 3.**
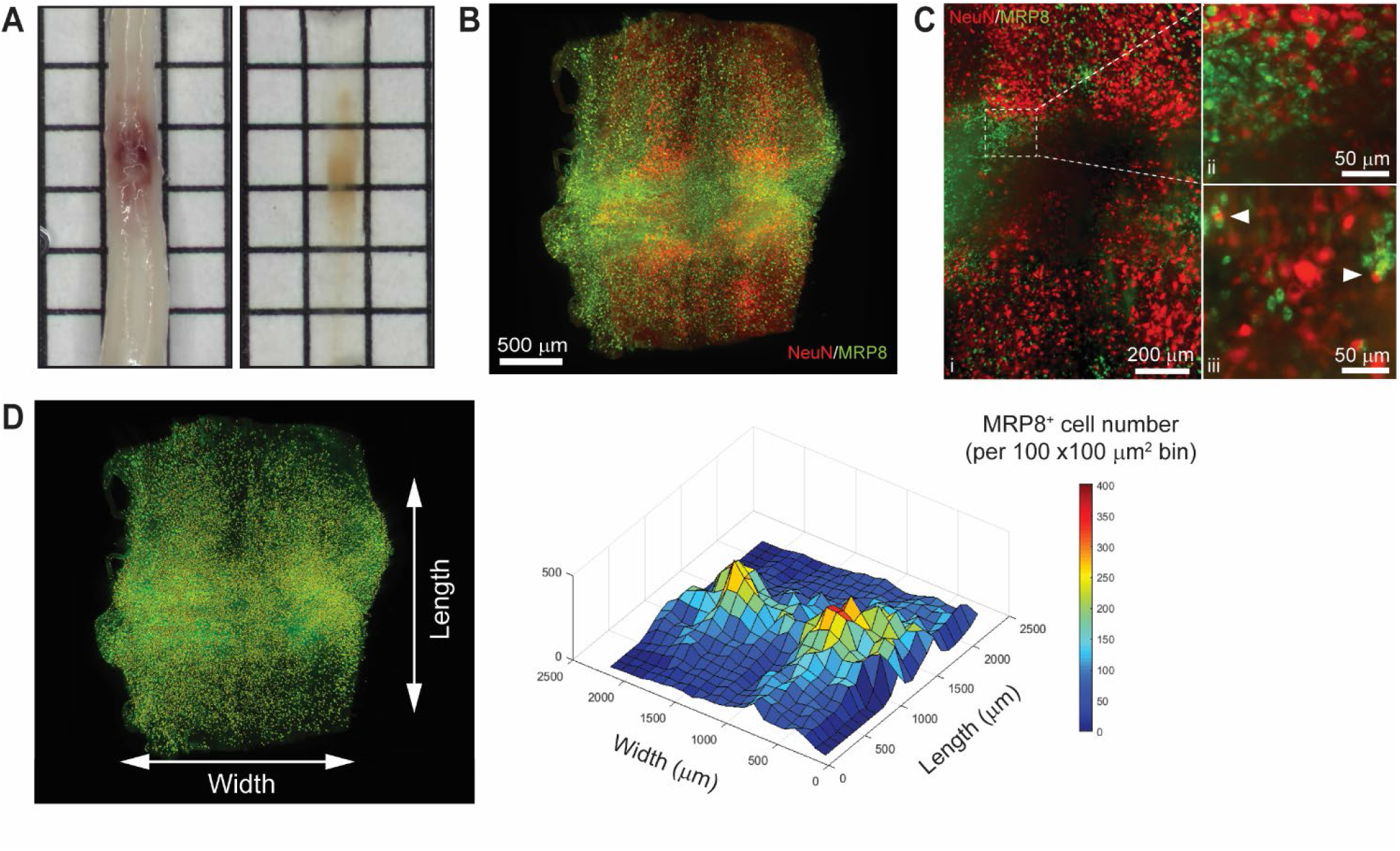
3D evaluation of inflammation in the acutely injured spinal cord using PACT clearing and lightsheet microscopy. A) PACT clearing of the acutely injured spinal cord. B) 3D projection of MRP8 and NeuN IF labeling in the cleared spinal cord at 1 day post-SCI. C) Maximum intensity projection of MRP8 and NeuN IF labeling at the SCI site. Loss of NeuN^+^ neurons and MRP8^+^ cells (degranulated neutrophils) is observed towards the lesion epicenter (i). Single z-plane images show MRP8^+^ cells interacting with NeuN^+^ neurons (ii; arrows). D) MRP8^+^ cell counting in Imaris (yellow dots) and heat map of the spatial distribution of MRP8^+^ cells near the spinal cord lesion. Abbreviations: spinal cord injury (SCI), immunofluorescent (IF).

The compatibility of immune cell surface markers with tissue clearing has yet to be shown, however, we identified MRP8 as a granular protein in neutrophils that could be readily labeled by IF in cleared tissues (Figure 3B-C). Neutrophils are the first circulating immune cell population to invade the spinal cord in large numbers after injury and numerous MRP8^+^ cells were observed throughout the injured spinal cord at 1 day post-SCI, demonstrating significant accumulation of neutrophils. Co-labeling with NeuN demonstrated substantial loss of neurons at the lesion epicenter. The loss of NeuN coincided with a loss in MRP8 labeling near the lesion epicenter, which may be due to neutrophil degranulation (Figure 3Ci). In addition, we observed MRP8^+^ cells directly interacting with NeuN^+^ neurons in single z-stack plane images (Figure 3Cii, arrows). Using Imaris software, we quantified the number of infiltrated MRP8^+^ cells and found a greater number of MRP8^+^ cells at the lateral edges of the SCI lesion (Figure 3D). These results demonstrate the potential for PACT clearing and 3D imaging to investigate the spatial dynamics of inflammation after SCI, which could provide critical insight for future therapeutic strategies such as immunomodulation.

### 4.4 Uniform labeling of white matter after SCI in PACT cleared tissues

One major limitation of utilizing whole tissues such as the brain and spinal cord is non-uniform labeling of abundant antigens due to outside-in labeling. A recent method to provide uniform labeling by controlling label interaction kinetics, called SWITCH, has been recently described to enable uniform labeling of myelinated structures within the mouse brain (Murray et al. 2015). To determine if SWITCH could be used to visualize white matter in the mouse spinal, as well as damage to the white matter during SCI, we performed SWITCH mediated labeling of myelin with DiD. First, we assessed clearing of injured spinal cords at 42 days post-SCI. Unlike the acutely injured spinal cord, the lesion site in the subacute phase of SCI could not be successfully cleared using PACT (Figure 4A, yellow arrows). However, the spared tissue region immediately surrounding the lesion did clear effectively and therefore could be readily imaged by lightsheet microscopy.

**Figure 4.**
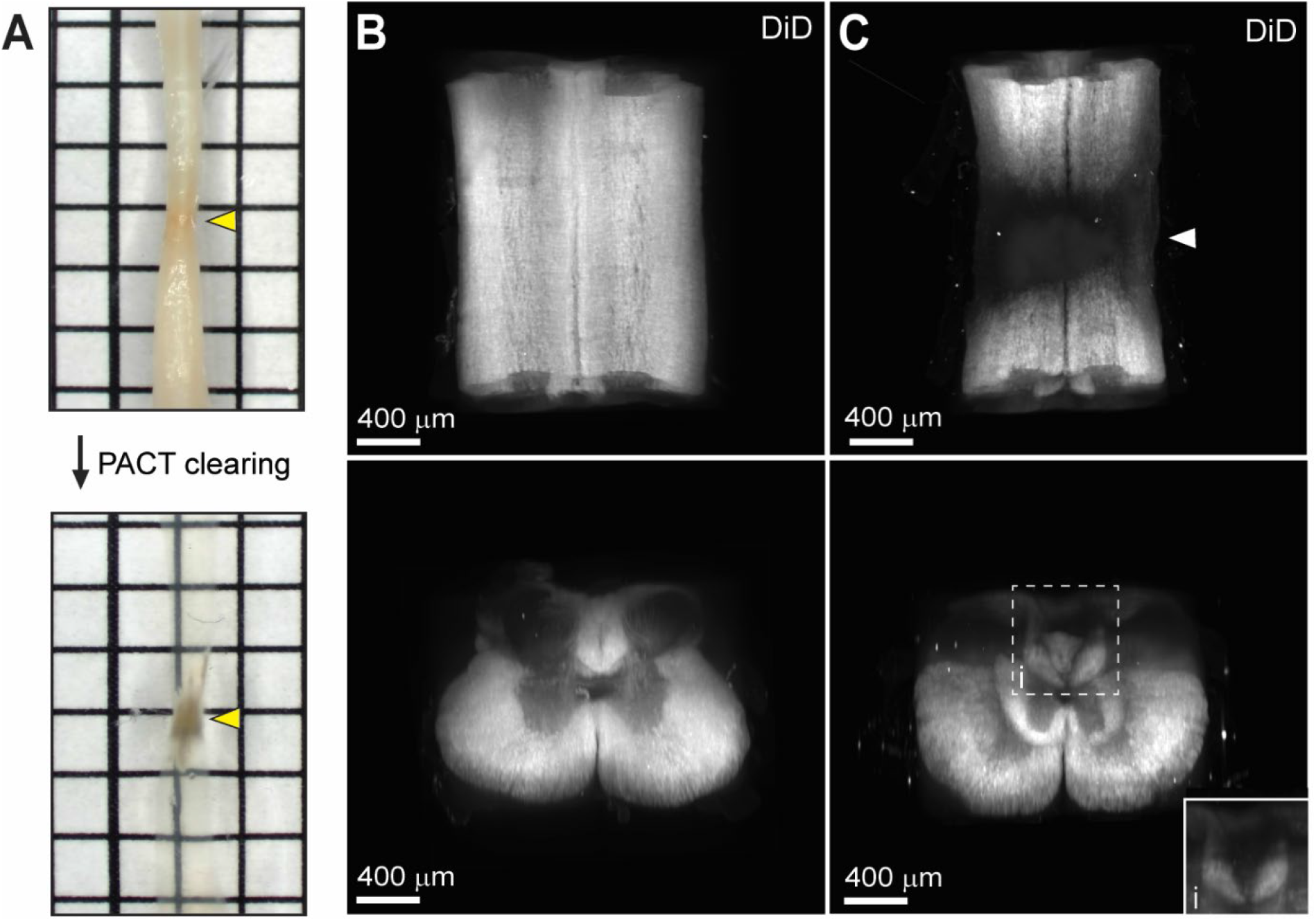
Uniform white matter labeling following clearing of the long-term injured spinal cord. A) PACT clearing of two injured spinal cords at 42 days post-SCI. Poor clearing of the SCI lesion (yellow arrows) does not prevent 3D imaging of the cleared surrounding tissue. B) 3D projection of SWITCH-mediated DiD labeling of the intact spinal cord from the dorsal and transverse views, respectively. C) 3D projection of SWITCH-mediated DiD labeling of the injured spinal cord (at 42 days post-SCI) from the dorsal and transverse views, respectively. Spared white matter tracks can be visualized around the SCI lesion site (white arrow). Abbreviations: spinal cord injury (SCI).

SWITCH-mediated labeling with DiD uniformly stained the white matter in the intact spinal cord (Figure 4B, Movie 3). While substantial loss of white matter labeling was observed around the SCI lesion site, spared white matter tracts could still be observed around the SCI lesion (Figure 4C, white arrow; Movie 4). Furthermore, loss of white matter labeling in the dorsal column was also observed rostral to the SCI lesion, which could be associated with specific regions of demyelination or dieback of damaged axons (Figure 4Ci). These data demonstrate a straightforward method for labeling myelin in the intact and injured spinal cord after PACT clearing.

### 4.5 PACT facilitates 3D visualization of transplanted neurons within the intact and injured spinal cord

Cell transplantation represents a promising solution to replace neuronal populations lost due to spinal cord injury (Fischer, Dulin, and Lane 2020). However, assessment of neuronal survival and integration has been limited to histological tissue sections that capture only a small fraction of the transplanted cell population. To determine if PACT clearing and lightsheet imaging could better capture the transplant cell population, we transplanted hiPSC-derived V2a interneurons at four injection sites in the intact and injured spinal cord and assessed their survival and axon extension two weeks later. In tissue sections, we were only able to observe cells labeled with a human specific Tau antibody (hTau) from one of the injection sites and long distance hTau ^+^ axon extension was difficult to assess due to axons entering and exiting the plane of the section (Figure 5A).

**Figure 5.**
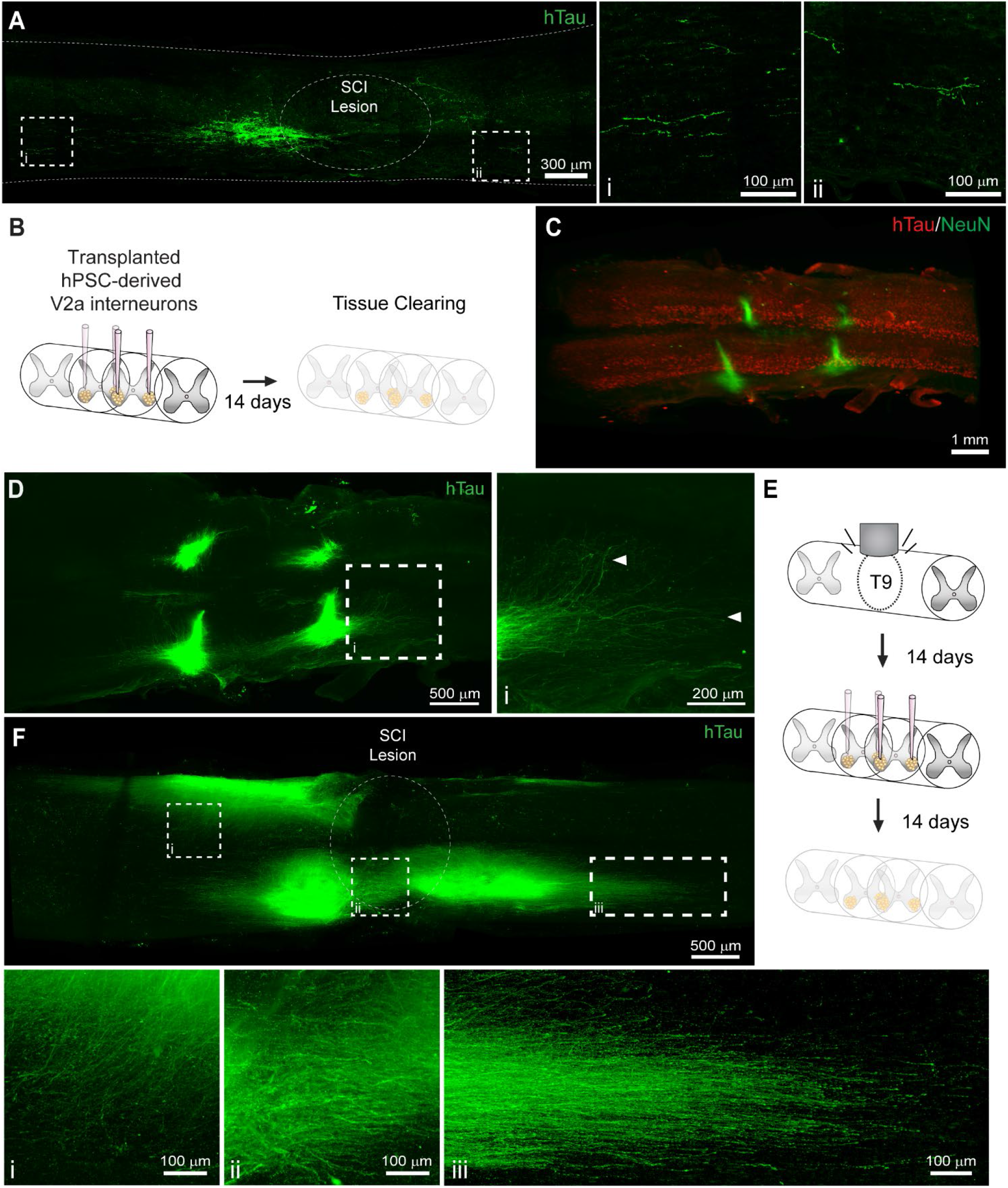
Labeling and 3D imaging of transplanted hiPSC-derived V2a interneurons in the cleared intact and injured spinal cord. A) hTau IF labeling in a sagittal section of the injured spinal cord at 14 days following transplantation. (i-ii) hTau+ axons can only be visualized for a limited distance. B) Schematic of hiPSC-derived V2a interneuron transplantation in the intact spinal cord and PACT clearing 14 days later. C) 3D projection of hTau and NeuN IF labeling in the cleared intact spinal cord showing transplanted hiPSC-derived V2a interneurons. D) Maximum intensity projection showing hTau+ axon extension in the host spinal cord. (i) In cleared spinal cords, hTau+ axons can be visualized for long distances from the transplant site. E) Schematic of SCI, hiPSC-derived V2a interneuron transplantation, and PACT clearing. F) Maximum intensity projection of hTau IF labeling of hiPSC-derived V2a interneurons at 14 days following transplantation in the injured spinal cord. Axon extension is observed into the grey matter (i), around the SCI lesion site (ii), and distally along the white matter (iii). Abbreviations: human induced pluripotent stem cell (hiPSC), human Tau (hTau), spinal cord injury (SCI).

Conversely, clearing the whole spinal cord and performing lightsheet imaging allowed us to easily visualize all four injection sites in the intact spinal cord by hTau ^+^ IF labeling (Figure 5B-C). In addition, we could readily observe hTau ^+^ axon extension from the transplant site into the host spinal cord (Figure 5D, arrows; Movie 5). We also transplanted hiPSC-derived V2a interneurons at two weeks post-SCI and collected spinal cord tissues for clearing two weeks later (Figure 5E). We observed variable hTau ^+^ cell survival at the four injection sites likely due to the less favorable environment of the injured spinal cord (Figure 5F, Movie 6). From multiple transplant sites we observed hTau ^+^ axon extension into the spinal cord grey matter (Figure 5Fi), around the SCI lesion site (Figure 5Fii), and along the spinal cord white matter (Figure 5Fiii). These results show a new and promising approach for assessing transplant cell behavior in the mouse spinal cord.

## 5 Discussion

This study describes a gentle and efficient method to clear the murine spinal cord through optimization of the PACT tissue clearing procedure. Hydrophilic and hydrogel-based approaches provide several advantages for tissue clearing including stability of endogenous and transgenic fluorescent protein signals, greater preservation of epitopes, increased tissue permeability for IF labeling, and compatibility with other aqueous-based histological staining procedures (Ueda et al. 2020). We modified several critical steps in the PACT clearing process which enabled robust clearing of the mouse spinal cord in as little as 6 days. While electrophoresis may further expedite clearing, we chose a passive approach to avoid the risk of tissue damage associated with electrophoretic tissue clearing, as well as the need for specialized equipment (Kim et al. 2015). The simple and reproducible approach described here is advantageous for enabling the broad applicability of spinal cord tissue clearing for many research laboratories.

Poor IF labeling is often observed in the thick CNS tissues used in tissue clearing studies due to limited diffusion and potential denaturation or purging of epitopes (Murray et al. 2015). With the optimized PACT methodology, we achieved complete penetration of antibodies into the cleared spinal cord tissue in as little as two days. We identified robust labeling of cholinergic neurons (ChAT), neuronal soma (NeuN), neutrophils (MRP8), and transplanted human neurons (hTau).These data demonstrate the promising ability of the optimized PACT tissue clearing protocol and IF labeling for lightsheet imaging of whole spinal cords from mice. One major limitation we discovered in our study was the incomplete staining of highly abundant epitopes, which can deplete primary antibodies during outside-in staining (data not shown). Recent methodologies, such as SWITCH (Murray et al. 2015), have been developed to sub-maximally label epitopes in a uniform manner throughout thick cleared tissues, which represents a critical first step in addressing the issue of non-uniform labeling.

Many of the studies utilizing tissue clearing and 3D imaging to study neural circuits in the spinal cord have relied on viral vectors encoding for fluorescent proteins such as enhanced green fluorescent protein (eGFP) and tdTomato to label specific neuronal populations (Asboth et al. 2018; Hilton et al. 2019; Ni et al. 2014; Soderblom et al. 2015; Wang et al. 2018). One limitation to these strategies is the potential quenching of fluorescent protein signals by tissue clearing procedures, especially with hydrophobic protocols such as 3DISCO and BABB (Soderblom et al. 2015; Ueda et al. 2020). However, we demonstrated that PACT clearing is compatible with transgenic tdTomato fluorescence. The fluorescent intensity of tdTomato in cleared spinal cords form Chx10::tdTomato mice was sufficient to visualize soma morphology and axon extension from V2a interneurons. This technique may be useful for mapping of spinal cord neural circuits and could be coupled with stochastic reporter mouse lines, such as Brainbow (Cai et al. 2013), to map dense neuronal populations across a broad fluorescent spectrum. Although not all fluorescent proteins are compatible with the PACT clearing process, common fluorophores are already being optimized for hydrogel-based tissue clearing applications (Scott et al. 2018). Combined with the optimized PACT clearing procedures presented, these advances can help elucidate the complex neural circuitry of the spinal cord.

Inflammation plays a critical role secondary injury following SCI, however, current techniques to assess immune cell populations, such as flow cytometry and histology, require dissociation or sectioning of the injured spinal cord tissue. At best, these techniques capture a limited view of the total spatial information. PACT clearing and IF labeling of MRP8 enabled easy visualization of non-degenerate neutrophils, which have not released their granule contents, throughout the entire cleared spinal cord. MRP8^+^ neutrophils were found to be most abundant near the lateral edges, but not the rostral and caudal edges, of the lesion at 1 day post-SCI. This striking distribution of neutrophils in the acutely injured spinal cord would be difficult to determine using 2D tissue sections, thus demonstrating the utility of tissue clearing and 3D imaging. One major limitation we found in our study was poor IF labeling with traditional immune cell markers in cleared tissues (data not shown). The incompatibility of PACT clearing, as well as other clearing techniques, with the cell surface markers commonly used to specify immune cell subtypes might be due to removal of the majority of cell membranes during the delipidation phase of clearing. One method to overcome this limitation is the use of fluorescent reporter mice with immune cell type specific Cre lines to constitutively label cells with fluorescent proteins, such as tdTomato, that are compatible with PACT clearing as described earlier. Future studies are needed to better understand the role of neutrophils in the acutely injured spinal cord, however, the data in our study demonstrate the potential for tissue clearing and 3D imaging to better elucidate neutrophil functions in the spinal cord parenchyma.

White matter sparing is a common labor-intensive analysis of residual tissue after SCI that is typically performed using transverse tissue sections that can be extrapolated into a low-resolution 3D models. We demonstrate that loss of white matter following SCI could be observed in 3D images of cleared spinal cord tissue at 6 weeks post-injury. The sub-chronic SCI lesion site failed to fully clear using the PACT protocol, however, similar results have been previously shown using hydrogel-based clearing techniques (Asboth et al. 2018). This may be due in part to the inability of hydrophilic clearing techniques to clear extracellular matrix rich tissues, as previously shown (Azaripour et al. 2016). To visualize residual myelin, we utilized a lipid dye-based (DiD) strategy previously developed with the SWITCH procedures for uniform labeling (Murray et al. 2015). White matter labeled with DiD was easily visualized in both intact and injured spinal cord tissue and spared white matter tracts could be readily observed outside of the spinal cord lesion. Further development of this approach could increase the accuracy and reduce the time necessary for white matter sparing analyses.

Cell replacement therapies are a promising approach to repair the damaged neural circuitry of the spinal cord (Butts et al. 2017; Fischer, Dulin, and Lane 2020), however, current histological procedures provide an incomplete interrogation of transplanted neurons in the host spinal cord. PACT clearing and 3D imaging enabled complete visualization of axon extension from transplanted hiPSC-derived V2a interneurons into the intact and injured spinal cord of mice. We identified transplanted human cells by IF labeling for a human specific form of the axonal marker Tau. As such, the described methods are limited to studying human neurons and more specifically their axons. However, fluorescent labeling of the transplanted cells could enable analysis of a wide variety of therapeutic cell populations from many different species. Our proof-of-concept studies demonstrate that tissue clearing and 3D imaging provide powerful insight into the assimilation of transplanted cell populations into the host spinal cord. The ability to interrogate the transplant architecture in cleared spinal cords may help improve future cell replacement strategies.

This report provides a detailed protocol for passive clearing of the mouse spinal cord that can be achieved without the need for specialized equipment. Fluorescent signals, either transgenic or from IF labeling, can be readily visualized in cleared tissues using confocal and lightsheet imaging. The new methodology has the potential to improve our understanding of spinal cord neural circuitry, mechanisms of secondary injury, and the discovery of effective therapeutic strategies. Future studies are needed to expand this technique to other species used for SCI research including rats and non-human primates.

## 7 Author Contributions

*“DM, NI, SS, and SC and TM contributed to conception and design of the study. RM, EM, JB, FJ, MP, HM, MK, and SC contributed to experimental design, data collection, and data analysis. DM wrote the first draft of the manuscript. FJ, MP, HM, and MK wrote sections of the manuscript. All authors contributed to manuscript revision, read, and approved the submitted version”*.

## 9 Acknowledgments

Use of the Texas A&M Microscopy and Imaging Center and the Gladstone Institutes Histology and Light Microscopy Core is acknowledged.

## 10 Data Availability Statement

All data is available from the authors upon request.

